# Global, Highly Specific and Fast Filtering of Alignment Seeds

**DOI:** 10.1101/2020.05.01.072520

**Authors:** Matthis Ebel, Giovanna Migliorelli, Mario Stanke

**Affiliations:** Institute for Mathematics and Computer Science, University of Greifswald, Walther-Rathenau-Str. 47, 17489 Greifswald, Germany; Center for Functional Genomics of Microbes, University of Greifswald, University of Greifswald, Felix-Hausdorff-Str. 8, 17489 Greifswald, Germany

## Abstract

An important initial phase of arguably most homology search and alignment methods such as required for genome alignments is *seed finding*. The seed finding step is crucial to curb the runtime as potential alignments are restricted to and *anchored at* the sequence position pairs that constitute the seed. To identify seeds, it is good practice to use sets of spaced seed patterns, a method that *locally* compares two sequences and requires exact matches at certain positions only.

We introduce a new method for filtering alignment seeds that we call *geometric hashing*. Geometric hashing achieves a high specificity by combining non-local information from different seeds using a simple hash function that only requires a constant and small amount of additional time per spaced seed. Geometric hashing was tested on the task of finding homologous positions in the coding regions of human and mouse genome sequences. Thereby, the number of false positives was decreased about million-fold over sets of spaced seeds while maintaining a very high sensitivity.

An additional geometric hashing filtering phase could improve the run-time, accuracy or both of programs for various homology-search-and-align tasks.

## Background

Aligning two or more genomic (or protein) sequences is one of the most fundamental tasks in bioinformatics. A base assumption is that if two sequences align well, they are likely to share a common evolutionary origin, i.e. are homologs. Often, one is especially interested in finding *orthologies* which indicate the same function. The alignment of whole-genomes is instrumental to comparative genomics and comparative genome annotation in particular [1]. The number of newly sequenced genomes can be expected to continue to grow for a long time. For example, the Vertebrate Genomes Project aims to generate reference genome assemblies of about 70,000 vertebrate species [2]. Creating a whole-genome (multiple) alignment requires to construct many local alignments of evolutionary related fragments of the different genomes. The task to find homologous genomic regions is of increasing importance and accurate, efficient and scalable methods are needed [1]. Thereby, most truly homologous genomic regions (e.g. coding exons of orthologous genes) shall be found but the number of hits of unrelated regions shall be limited.

A common approach to alignment is the seed and extend approach [3]. In a first step, very short local similarities are sought. For the sake of speed, these similarities may be required to be identities. The resulting hits then serve as *alignment anchors*. In a second step, starting from these anchors, local alignments are computed. As the second step is usually more time consuming, it is important that the anchors, also called *seeds*, from the first step are sensitive and specific, while being found very quickly. As sensitiviy and specificity can be traded off against each other we will sometimes generally refer to accuracy. This work focusses on finding these seeds and improving established methods to do so.

In early aligners, small exact matches of length *k*, so called *k*-mers, were used as seeds [3]. Ma et al. [4] and Burkhardt and Kärkkäinen [5] were among the first to introduce the idea of *spaced seeds*, where bases at certain positions of a seed – so called *don’t care* positions – are not required to match. They indepentently found that the positions, at which mismatches are allowed, have a high influence on the sensitivity of spaced seed patterns. In a following article, Li et al. [6] introduced multiple spaced seeds and use a *set of spaced seed patterns* to find alignment anchors with even higher sensitivity. Much research has been carried out investigating the optimality of spaced seed patterns and the hardness to actually compute such patterns (e.g. [7, 8, 9, 10, 11, 12, 13, 14, 15, 16, 17, 18, 19, 20], among others), usually under the assumption of very simple probabilitic models of homologous and non-homologous sequence pairs. The interested reader is referred to this survey by Brown [21] for a more detailed overview of the topic.

Spaced seeds have been applied in alignment software such as DIAMOND [22], LASTZ [23], YASS [24] and the discontiguous MegaBLAST version of BLASTn [25, 26, 3], to name a few. In previous research on further improving spaced seeds, Noé and Kucherov [27] introduced an additional filter criterion, requiring an anchor to have neighbouring seed matches on close diagonals. Mak et al. [28] presented “indel seeds” that can cope with very small indels in homologous regions, thereby sacrificing speed. Recently, Leimeister et al. [29] applied spaced seeds with an additional filtering step in a multiple sequence alignment pipeline. They used very sparse spaced seed patterns with 10 match and 100 don’t care positions and a novel filtering step, scoring *all* positions in a seed to filter out noise.

In this article, we compare fast methods to increase sensitivity and specificity of (spaced) seeds. We here consider methods that are based on exact re-occurrences of fixed-length sequence patterns, which can be implemented very efficiently and thus can be used as a very fast initial step in an alignment pipeline. Typically, such seeds could be in a particularly well-conserved region with higher sequence similarity and would be used as starting point for a sequence of steps that constructs a local alignment that extends further and uses a detailed scoring to increase the specificity of the hit. Subsequent filtering steps can be slower than the seed finding and typically include a gapless extension of anchors (e.g. with the X drop heuristic [3]), thresholding the alignment score of the extension, the joining of nearby gap free alignments and local alignment extensions that may contain gaps and further thresholding ([3, 23]). The seeds can be interpreted as *anchors* that constrain the set of admissible alignments, significantly reduce their number and also the runtime of alignment algorithms.

We introduce a novel method, *geometric hashing*. Geometric hashing is a fast filter of candidate seeds taken, such as exact *k*-mer matches induced by spaced seed patterns. To achieve a higher accuracy, the matches from homologous regions are accumulated over possibly *long distances* using a secondary hashing technique. We evaluate the methods on real genomic data from human and mouse for sensitivity, as well as on artificial random sequences to assess specificity. Geometric hashing can be adjusted to simultaneously be more sensitive and much more specific than existing methods at finding seeds in coding regions of homologous genome regions and requires only a small fraction of additional runtime.

We confirm that multiple spaced seed patterns are better than a single spaced seed pattern. On this task, sets of four spaced seeds produce one to two orders of magnitude fewer false hits than a single spaced seed pattern. We also confirm and quantify the superiority of spaced seeds over contiguous *k*-mers as seeds in finding homologous exons.

*Geometric hashing* can be adjusted to decrease the number of false positives by at least six orders of magnitudes while only marginally decreasing the sensitivity from 97.4% to 97.2%. Alternatively, when using a *k* that is one smaller than the *k* used by a set of four spaced seed patterns, here *k* = 14 versus *k* = 15, geometric hashing simultaneously reduces the false negatives by 19% from 2.6% to 2.1% and the number of false positives by a factor of about 3 · 10^5^.

## 1 Methods

### 1.1 Test Data

As test data set we used genomic sequences from the softmasked genome assemblies of human and mouse:

- *Homo sapiens* (hg38 GCA_000001405) [30]
- *Mus musculus* (mm10 GCA_000001635) [31]

Ortholog protein coding genes have been queried from Biomart [32], the Ensembl interface to access homology predictions of genes. Pairs of ortholog genes were selected, thereby ensuring a high confidence in the orthology relation according to their respective Ensembl score [33]. Further, only one-to-one orthologs were allowed in the dataset, such that each sequence appears in only one pair and no two sequences from different pairs were considered orthologs.

We retrieved the `GFF` genome coordinate files containing Ensembl annotations for the human genome, a single representative principal transcript was picked for each gene in order to avoid any bias towards those genes which include a larger number of transcripts.

We selected a subset of 705 pairs of human and mouse gene regions, each containing one orthologous gene pair. The average sequence lengths were 74 *kbp* and 65 *kbp* for human and mouse, respectively. The total length of human and mouse genic regions in the test set were approximately 52 *Mbp* and 46 *Mbp*, respectively.

As real genome sequences cannot be guaranteed to be void of further homologies besides the chosen orthologies, we simulated a set of random sequences for an estimation of the number of false positives. For each real sequence in the human-mouse dataset, we simulated a random DNA sequence with independent and uniformly distributed nucleotides of the same length as the respective gene, labeling it with the respective genome. Choosing this simple distribution for the negative examples is in agreement with most previous work on spaced seeds. The independent and uniform distribution is stated either explicitly [34, 21] or is implied by considering all seeds of length or weight *k* as equally specific [35, 11]. All seed hits in this artificial set of DNA sequences were counted as false positives.

### 1.2 Evaluation

As spaced seeds have generally been designed to anchor an alignment of *two* sequences, we will evaluate and compare all methods on a *pair* of genomes as well, here from human and mouse. However, the geometric hashing idea generalizes to more than two genomes. Formally, we define an alignment seed as a quadruple (*S*_1_, *i, S*_2_, *j*), where *i* is a position in sequence *S*_1_ and *j* is a position in sequence *S*_2_. The seed can be interpreted as a prediction that these two positions are believed to be homologous positions. The applied methods actually rather identify small region pairs of equal length, e.g. of length *k*. We therefore use for evaluation purposes the region midpoints.

Several alignment anchors could eventually lead to the same local alignment of homologous regions. This is to be expected for seeds (*S*_1_, *i, S*_2_, *j*) and (*S*_1_, *i*′, *S*_2_, *j*′) of the same sequence pair, where *i*′ − *i* and *j*′ − *j* are small and similar or even equal. It is therefore sufficient to find at least one of the anchors that are redundant in this sense, which motivates the following accuracy measure.

#### 1.2.1 Sensitivity

We say a seed (*S*_1_, *i, S*_2_, *j*) *supports* a coding sequence (CDS) with coordinate range [*a, b*] of sequence *S*_1_ if *a* ≤ *i* ≤ *b* and if (*S*_1_, *S*_2_) is a pair of homologous gene regions. We calculate the percentage of human coding exons (CDS) with *at least one* supporting seed and define

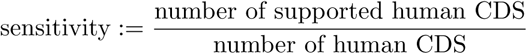

This measure is based on the notion that the alignments of coding sequences of homologous CDS pairs typically contain only small numbers of indels, are relatively highly conserved and therefore a subsequent alignment anchored in such a seed would likely result in an alignment of at least a large part of the exons. Brejova, Brown and Vinar find on human-mouse homologous coding gene pairs that the CDS alignment fragments, that are not interrupted by introns, are on average 152 bp long. This figure reduces to 120 bp if the fragments are considered to end at an indel [10].

Note that we did not require that the corresponding position *j* in mouse is homolog. One reason for this choice is that the accuracy measure would otherwise depend on the completeness and correctness of some reference alignment and would also presuppose that a matching splice form is annotated and identified. The sensitivity would then have an unknown upper limit < 1 that depends on other tools and their settings. Another reason is that we will below consider limits to the overall number of false positives such that the number of false positive seeds in homologous region pairs (*S*_1_, *S*_2_), that contain only a single gene each, is negligible. Thirdly, all methods are compared with the same accuracy measures and we do not expect the relative performances to be effected.

Note that this sensitivity measure also cannot quite be expected to achieve 100% because not all human exons have a homologous region in the mouse genome. Nevertheless, we consider choices of *k* and other parameters most relevant when the sensitivity is at least 0.9.

#### 1.2.2 False Positives

To measure and compare the prediction of wrong seeds we applied all approaches also to the set of random sequences described in Section 1.1. Any hit between a random ‘human’ and ‘mouse’ sequence is considered a false positive (FP). We normalize the number of false positives #FP to 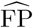 as follows

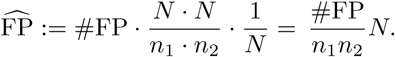

#FP is the total number of counted false positives. *n*_1_ = 51.878 · 10^6^ and *n*_2_ = 45.741 · 10^6^ are the total lengths of the random ‘human’ and ‘mouse’ sequences from the artificial data set. *N* = 3.22 · 10^9^ is the size of the human genome. 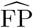 can be interpreted as the extrapolated fraction of false positive seeds per genome position if two human-sized genomes were compared.

The rationale behind this measure is as follows. The alignment space – more partiular, the set of all admissible (*S*_1_, *i, S*_2_, *j*) – is inherently of quadratic size *N* ^2^ for two genomes of total size *N*. However, through appropriately large *k*’s or thresholds, seed-finding can reduce the number of hits that are further examined to something that is linear in the genome size(s) *N*. An effort that is linear in *N* is unavoidable and acceptable.

### 1.3 Seeding Approaches

In this section five methods M1-5 for seed finding are described in order of increasing accuracy and sophistication. M1 searches for contiguous matches, M2 and M3 use a single and multiple spaced seed patterns, respectively. Methods M4 and M5 use seed candidates found by M3 and apply additional filtering steps to reduce 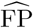.

Matches found by M1-M3 are small similar region pairs, thus we extract the respective midpoints as seed candidates. Repeats in the sequences could lead to many seed candidates originating from the same *k*-mers. Firstly, we do not consider sequence positions that fall in a repeat-masked part of the genome. Secondly, we apply a simple filter to reduce this noise for all methods. If a *k*-mer leads to more than ten seed candidates, we only compute a random subsample of size 10 of all possible seed candidates. Ignoring seeds of patterns with many matches is in accordance with filtering techniques used in other genome aligners, e.g. DIAMOND [22].

For a string *S* and sequence positions *a* ≤ *b* let *S*[*a..b*] denote the substring of *S* from position *a* up to and including position *b*. Exclusion of the end position is denoted with round parentheses, e.g. *S*[*a..b*) goes up to position *b* − 1 only. In the following, all sequence positions are implicitly assumed to be in the range of the sequence length. Seed matches with ambiguous or unknown characters (e.g., `n`) were not considered a match and were discarded.

#### 1.3.1 M1: Exact Contiguous Matching

This simplest method serves as a baseline. For a given weight *k*, we say that two sequences *S*_1_ and *S*_2_ have an exact contiguous match at position pair (*a, b*) if and only if

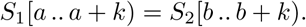

We then take (*S*_1_, *a* + ⌈*k/*2⌉, *S*_2_, *b* + ⌈*k/*2⌉) as the *seed*, i.e. the center from the two identical substrings of length *k*.

#### 1.3.2 M2: Spaced Seeds

A spaced seed pattern is a binary pattern *p* = (*p*_1_, …, *p*_*ℓ*_) {0, 1}^*ℓ*^ of length *ℓ*, i.e. a string over the alphabet 0, 1 where the 1’s are called *match* positions and the 0’s are *don’t care* or *wildcard* positions. The length *ℓ* is called *span* and the number of match positions is its *weight k*, with *k* < *ℓ*.

The *k*-mer *x*_*i*_ is the string induced by applying *p* to a sequence *S* at some position *i*, concatenating only the characters from *S* that pair with a match position. More formally, let

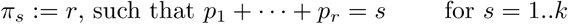

be the position of the *s*-th 1 in *p*. The *k*-mer *x*_*i*_ defined by

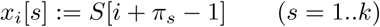

is said to be the *contiguous pattern induced* by spaced seed pattern *p* at position *i* in *S*.

Let *x*_*i*_ and *y*_*j*_ be the *k*-mers induced by a spaced seed pattern *p* of weight *k* at positions *i* − ⌈*ℓ/*2⌉ − 1 in *S*_1_ and *j* − ⌈*ℓ/*2⌉ − 1 in *S*_2_, respectively. We then say that *S*_1_ and *S*_2_ have a match according to the spaced seed pattern *p* at position pair (*i, j*) iff *x*_*i*_ = *y*_*j*_. Note that in the literature about spaced seeds, the notion is often used to refer to the binary spaced seed *pattern*. In this article, we call a seed a pair of sequence positions (*S*_1_, *i, S*_2_, *j*) that could serve as an alignment anchor and write “spaced seed *pattern*” when we refer to the binary pattern of *match* and *don’t care* positions.

It is well-established that the choice of *p* has a significant influence on the sensitivity even at a fixed weight [4, 9]. The sensitivity of spaced seed patterns is related to the number of overlapping hits [36]. Hits of contiguous seed patterns (M1) tend to cluster, while hits of seed patterns with low self-overlap are more evenly distributed and thus more sensitive [11]. Thus, one needs to use optimized spaced seed patterns to get the best results. We used the software `SpEED` [17, 35] to compute spaced seed patterns of desired weight, that are good under its simplifing assumptions on the distribution of homologous sequences. The underlying model requires the length of the homologous region and the similarity of the homologous region for which the seeds should be optimal. As in [4] we set the region length to 64. The base pair match probability was set to 0.85, the percent identity of corresponding CDS in human and mouse [37].

#### 1.3.3 M3: Set of Spaced Seed Patterns

Let *P* be a set of *m >* 1 spaced seeds patterns, each of weight *k*. We say that *S*_1_ and *S*_2_ have a match according to a spaced seed pattern *p* at position pair (*i, j*) if the two sequences have a match at this position pair for *any spaced seed pattern p* ∈ *P*. Li et al. introduced this concept to increase the sensitivity of spaced seed patterns. While reducing the weight *k* of a single spaced seed pattern also leads to higher sensitivity, the trade-off with getting more random hits at the same time is better when using more spaced seed patterns instead [4]. We again used `SpEED` to generate good sets of spaced seed patterns of desired weight with the same parameters as above.

These first three methods are well known and used as basis and baseline for the upcoming methods. The following methods describe additional filtering steps which can be applied to sets of seeds found by either of the former methods.

#### 1.3.4 M4: Neighbouring Matches

After all matches have been identified, locally the number of consistent matches are counted and a hit is only reported if the number of neighbouring matches reaches a threshold *τ >* 1. The idea is to allow the individual seeds to be less specific (smaller weight *k* or higher number of patterns *m*). Applying this filter, an isolated match that could be random and non-homologous does not necessarily result in a false hit. This method is similar to the one described by Noé and Kucherov [24].

What is considered local versus non-local is controlled by a variable *D*. Suppose (*S*_1_, *i, S*_2_, *j*) is a *candidate* seed from method M3. This position pair is reported as hit *only* if the total number of seeds of some positions (*S*_1_, *i*′) and (*S*_2_, *j*′), such that *i*′ − *j*′ = *i* − *j* and |*i* − *i*′| ≤ *D/*2, is at least *τ*. As customary, we call the sets {(*i, j*) | *i* − *j* = const} *diagonals* in the pairwise alignment space. In other words, all matches are counted in the region pair of length *D* centered around *I* and *j* that are on the same diagonal. Summarizing nearby hits on the same diagonal can be done very efficiently (see Section 1.4). As reporting hits on the same diagonal that are very close to each other are likely to be redundant, we do not allow two matches (*i, j*) and (*i*′, *j*′) to overlap, i.e. |*i* − *i*′| ≥ *ℓ*.

#### 1.3.5 M5: Geometric Hashing

Methods M1-M3 only consider the directly matching regions. Method M4 considers the immediate neighborhood only. In contrast, M5 is able to gather evidence for homology from matches that are distant to each other. The idea is to collect seed candidates from multiple exons from the same orthologous genes. Or, more generally, from seed candidates with similar distance differences in the two sequences. Reporting only seeds from sufficiently large such collections then increases specificity as single random matches typically remain unreported.

Our geometric hashing approach is motivated by an eponymous technique from object recognition in computer vision [38]. There, first distinctive points are identified in the image. A hash table is built, where the keys are ordered pairs from the distinctive points and the value is a collection of coordinates of the remaining points measured in a coordinates system given by the pair of key points. This is done for any two points from the object. To recognize an object in an image, the same is done for two arbitrarily chosen distinctive points. If the object is “known”, there will be a slot in the hash table that has a very similar collection of points relative to their key points, and thus the object can be predicted. Geometric hashing is robust to object rotation, translation, small variations in object shape and to partially occluded objects.

When transferring this concept to seed finding for pairwise sequence alignments, matters even get easier. The “objects” we try to identify are orthologous genes. Exons of orthologous genes are much better conserved than typical noncoding sequences, thus we expect many matches there, while there should be only few matches in the less conserved introns and intergenic region. The seeds are our distinctive points. The only transformation we require is horizontal shift. Two seed candidates (*S*_1_, *i, S*_2_, *j*) and (*S*_1_, *i*′, *S*_2_, *j*′) from different exons from the same two orthologous genes may be thousands or even ten thousands of basepairs apart in the genome. Yet, the relative distances *i* − *j* and *i*′ − *j*′ are often similar. Using this relative distance, we are able to collect seeds from even very distant exons. As the interjacent introns often have undergone length changes through insertions or deletions, we round the relative distances to multiples of *F* = 10, 000 *bp* (‘quantization’). This way, the relative distances of seeds become robust to varying intron lengths up to a certain degree. See Figure [1b)] where this is illustrated. It displays two sequences *S*_1_, *S*_2_ from human (top) and mouse (bottom), which turn out to be the orthologous glutathione synthetase (GSS) genes. The thick blue bars are annotated exons, which the algorithm is unaware of. The orange lines are seeds found by our geometric hashing approach. Some introns quite visibly differ in length, yet the seeds all were collected at one place we call a *tile*. A more formal description of the geometric hashing approach follows.

Let *S* be a set of candidate seeds, e.g. from either method of M1-M4. Let *F* be a *tile size* (we use *F* = 10, 000 for genomes) and define a *geometric map*

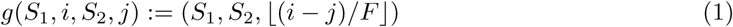

that maps a candidate seed to the pair of sequence identifiers and a difference tile. Here, ⌊⌋ means rounding down to the next integer. Let *T* := *g*(*S*) be the image set of the geometric map. We call the elements of *T tiles*. For a tile *t* ∈ *T*, the set *g*^−1^(*t*) contains seeds of the same sequence pair that have similar differences *i* − *j*:

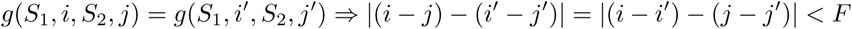

See Figure [1a)] for an illustration of the idea. One can think of a tile as a diagonal stripe of width *F* in the pairwise alignment space.

**Figure 1:**
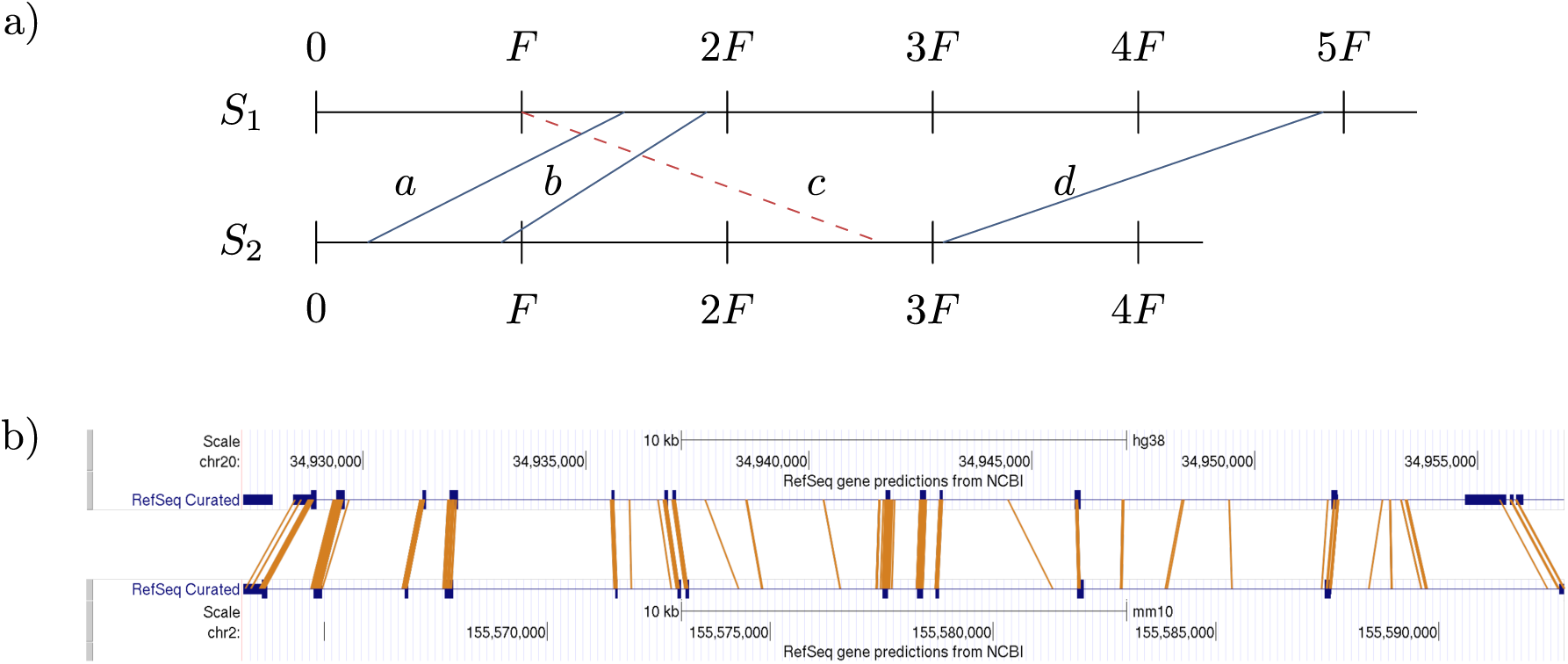
Geometric hashing. **a)** Idea of geometric hashing. Seeds *a, b* and *d* map geometrically to the same tile (*S*_1_, *S*_2_, 1) = *g*(*a*) = *g*(*b*) = *g*(*c*) and support each other even though they are distant and there are indels between them if they specify homologous site pairs. Seed candidate *c* maps geometrically to tile *g*(*c*) = (*S*_1_, *S*_2_, 2) whose significance falls below the threshold and is not reported as no other seeds map geometrically to the same tile. **b)** Seeds from human and mouse gene *glutathione synthetase* (*GSS*, Ensembl IDs `ENSG00000100983` and `ENSMUSG00000027610`, respectively). Conserved exons (thick blue bars) are hit by many seed matches (orange lines). All seeds from the ≈ 30 *kpb* gene range were collected in a single tile, despite differing intron lengths between corresponding exons. Edited screenshots from UCSC Genome Browser [39, 40]

The choice of a tile size in the order of 10, 000 *bp* was based on an analysis of our dataset (Figure [2]). The choice constitutes a tradeoff. Ideally, the tile size would be big enough so that even genes with greatly differing intron lengths are not ‘broken’ into neighbouring tiles. On the other hand, very large tile sizes may lead to spurious tiles that by chance contain many uninformative seed candidates.

**Figure 2:**
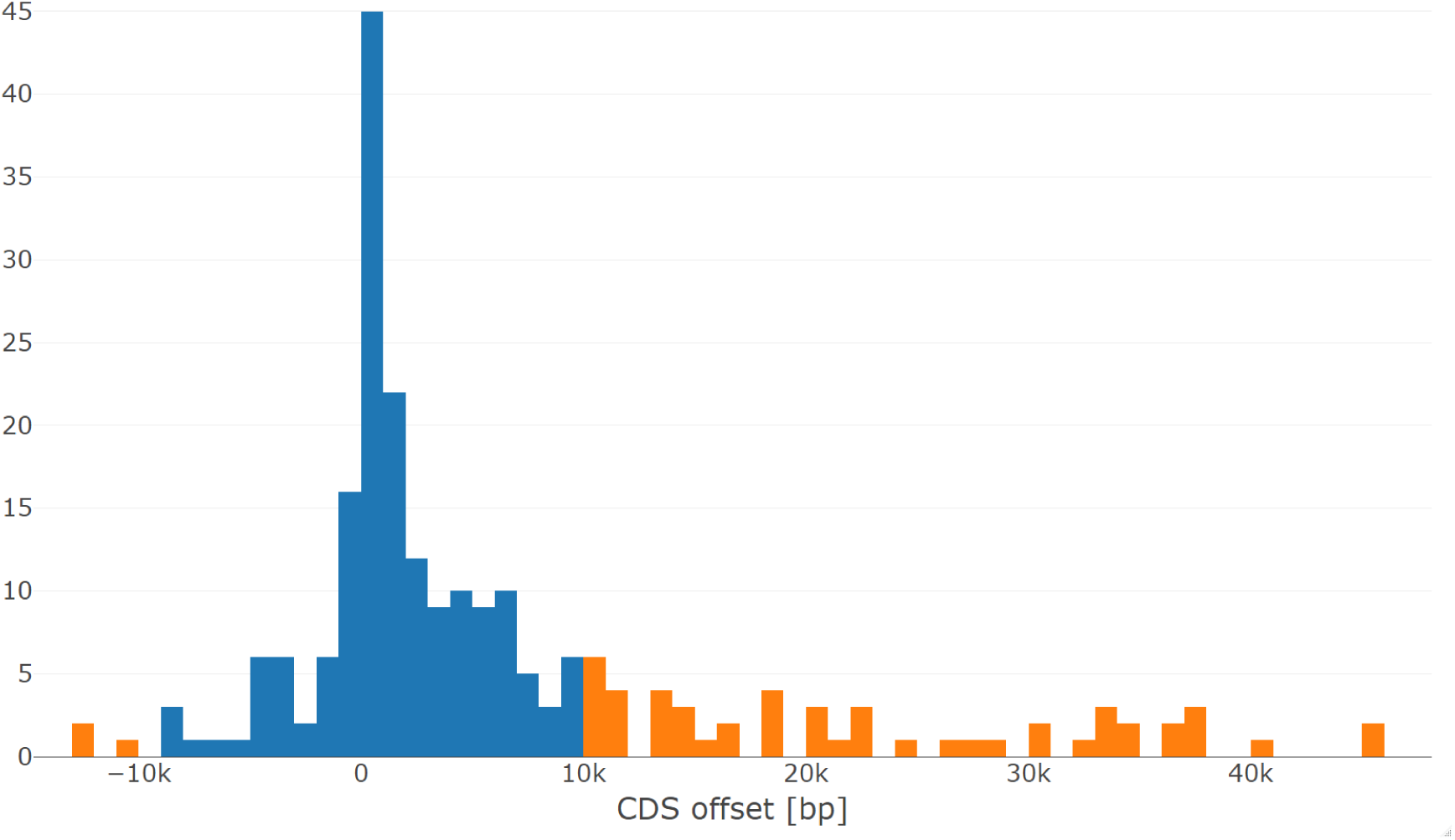
Distribution of maximal exon offsets Ω in the test set of human-mouse orthologs. Exon offsets measure the cumulative effect that indels have on the relative positions of exons. For most genes, maximal exon offsets are in the range [−10000 *bp*, 10000 *bp*] (blue). The distribution leans to the right, apparently because there are more transposable elements inserted in intronic regions in the human lineage. Outliers are not shown. See main text for the definition of Ω.

To study the effect of indels on the relative positions of exons of orthologous human-mouse pairs of genes, we filtered the human and mouse gene pairs from our data such that all pairs have the same respective number of CDS and assumed that identically numbered CDS are orthologous. We then calculated the maximum offset between ‘orthologous’ CDS: Say there are *n* CDS in a particular gene and denote the start of the *i*-th CDS in human with 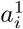 and in mouse with 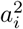. The maximum offset is then defined as 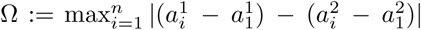. When looking at the distribution of maximum offsets (Figure [2]), we see that with a tilesize of 10000 *bp* we can catch approximately 75 % of genes in a single tile, while results show a reasonable 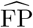.

After performing geometric hashing, the encountered tiles are scored and all seed candidates from a tile *t* are reported as seeds if the score of *t* matches or exceeds a threshold *τ*. If the threshold is not reached, none of the seed candidates of the tile are output. The tile scoring algorithm works as follows. Let |*S*_1_| and |*S*_2_| denote the lengths of sequences *S*_1_ and *S*_2_, respectively. We divide each tile into a fixed number of *b* sub-tiles of equal width *F/b*. Each seed candidate (*S*_1_, *i, S*_2_, *j*) ∈ *g*^−1^(*t*) in a tile *t* is assigned to a sub-tile ⌊*db/F*⌋ ∈ {0, …, *b*−1}, where *d* = *i*−*j* −*t*^∗^ is the seed candidate’s diagonal inside *t* and *t*^∗^ := *F* · ⌊ (*i* − *j*)*/F*⌋ is the first diagonal of tile *t*. We found that with our data, *b* = 500 sub-tiles work well. We denote with *n*_*r*_ the number of seed candidates in sub-tile *r* ∈ {0, …, *b* − 1} of *t* and sum the squares of all *n*_*r*_ for the score (Equation (2)). This gives a relatively higher score to tiles whose seed candidates cluster on few sub-tiles and less so for evenly scattered seed candidates. The former we expect in truly orthologous sequences where single orthologous exons share many seeds on similar diagonals (see Figure [1b)]), the latter we expect from random or unrelated sequences.

With longer sequences *S*_1_, *S*_2_, we expect more spurious seed candidates appearing by chance and normalize the score to account for this noise. We estimate the tile area *A*_*t*_, which is the maximal number of seed candidates (*S*_1_, *i, S*_2_, *j*) a tile *t* could theoretically have. The estimation formula is *A*_*t*_ := (*v* − *u*)*F*, where the terms *v* = *min*(|*S*_2_| + *t*^∗^, |*S*_1_|) and *u* = max(*t*^∗^, 0) are used to calculate the length of the starting diagonal of the tile. This estimation is reasonable near the main diagonal (i.e. *i* ≈ *j*) but not at the extreme cases where *i* ≫ *j* or *i* ≪ *j* and the true area is small. However, our experiments showed that it is not helpful to normalize scores of smaller tiles, so this is not an issue as we designed the normalization to only apply to large tiles. The scoring formula with the normalization term is

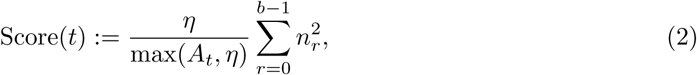

where *η* is a normalization parameter we set to *η* = 3 · 10^8^, which is roughly the median of all tile’s areas. Thus, for the smaller half of tiles the score is simply the sum of squared seed candidate counts in each sub-tile. For the larger tiles, this score is lowered according to the tile area.

### 1.4 Geometric Hashing Algorithm

We here outline the geometric hashing algorithm (M5) with pseudocode and provide a runtime analysis. The algorithm for M4 (Neighbouring Matches) is detailed in the supplementary material.

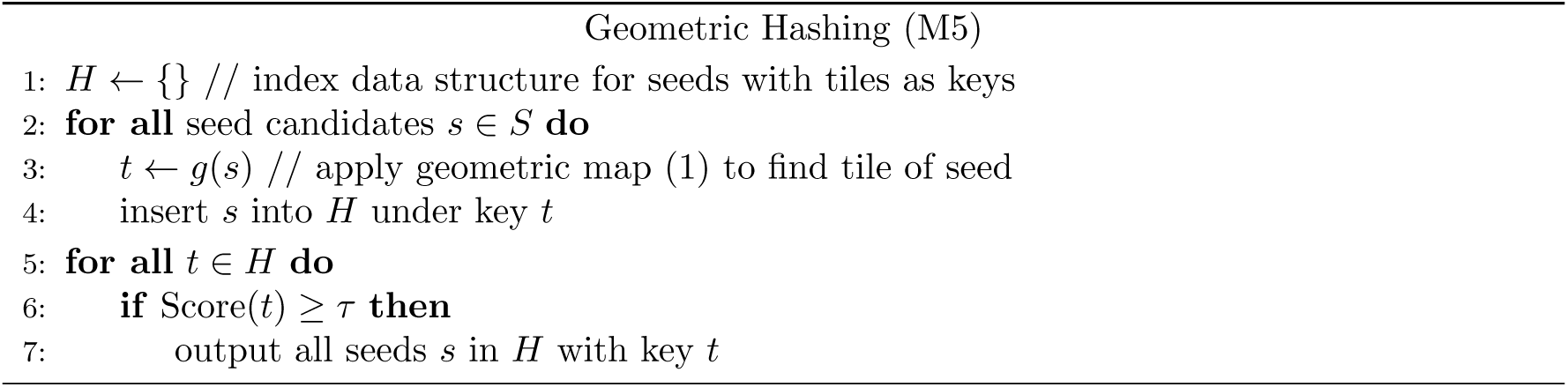

Our implementation in C++ uses a hash table for the data structure *H*. Inserting and querying elements from such a data structure can be done in *O*(1) expected time. The key of the hash table is a tile, and the value is an unordered set of seed candidates that fit into the tile. Inserting a seed candidate into an unordered set also takes *O*(1) time. Computing the score in line 6 is linear in the number of seed candidates in the tile.

## 2 Results

Figure [3] shows the extrapolated false positive fractions and sensitivities for all five methods, when the weight *k* and the number of spaced seed patterns (for M3) is varied. A decreasing *k* leads to larger sensitivities but the number of false positives grows exponentially. With a weight *k* = 13 all methods except M4 are expected to produce more than 10 times as many seeds or intermediate seed candidates (M5) as there are bases in one genome, when seeds in two human-sized genomes are searched. For such tasks, a further decrease of *k* could quickly render seed finding or seed extensions computationally infeasible or have prohibitive memory usage.

**Figure 3:**
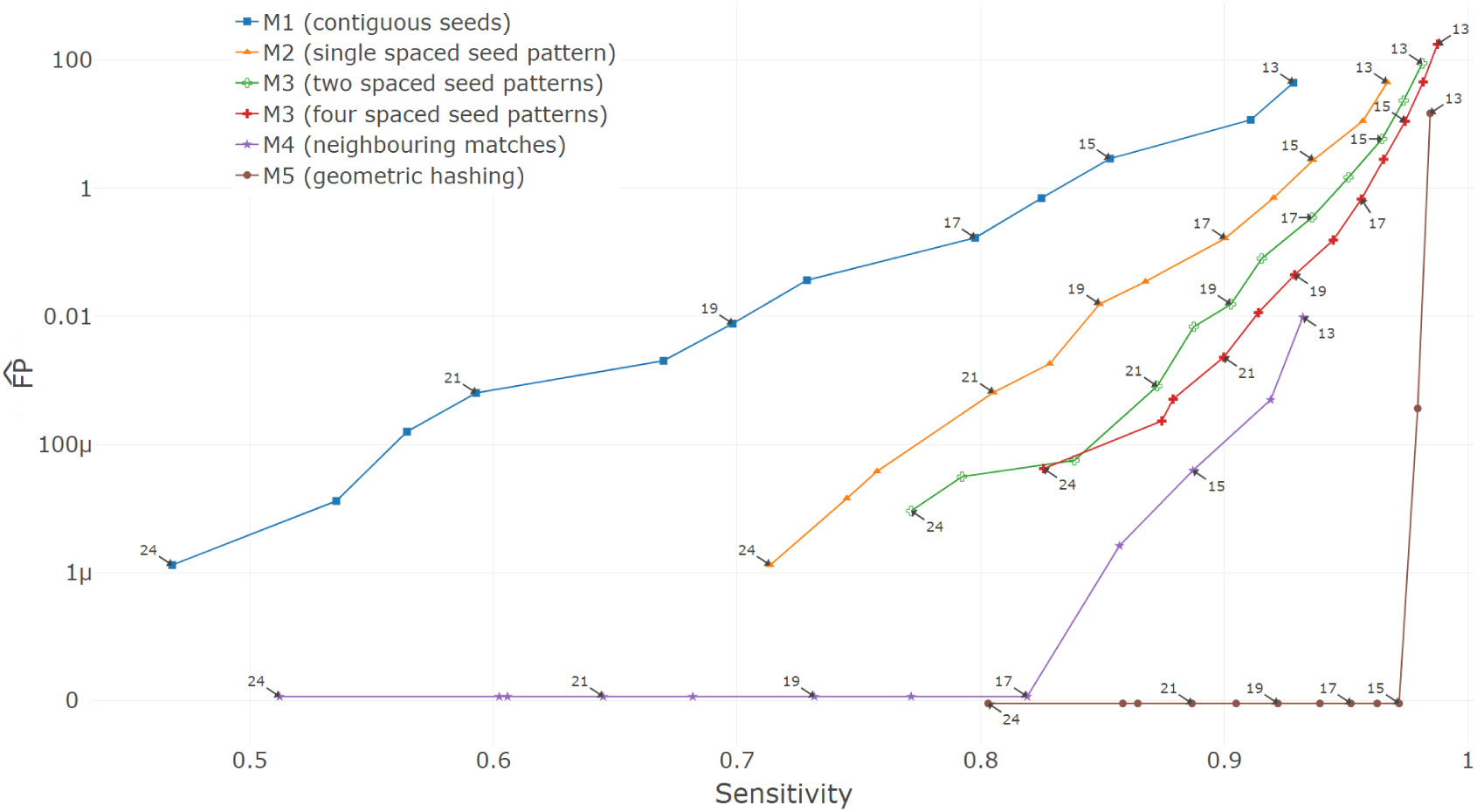
Comparison of the different methods. The *y*-axis shows on a logarithmic scale the total number of false positive seeds that are scaled to be estimates of the total number of false positive seeds per base in the genome if two complete human-sized genomes were compared (*µ* means 10^−6^). The *x*-axis is the percentage of coding exons that are supported by seeds. Each data point represents a run of the respective method with a certain weight *k* (point labels), ranging from 13 (top right points) to 24 (bottom left). Note that data points at 0 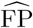 are slightly shifted for better visibility.

On our test data set of unrelated sequences, geometric hashing completely removed all false positive seeds for all *k* ≥ 15, while maintaining a very high accuracy (bottom right data point in Figure [3]). For methods that predicted 0 false positives in these random sequences (M4 for *k* ≥ 17 and M5 for *k* ≥ 15), the confidence interval for 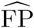 is [0, 6.2 · 10^−6^] (confidence level 1 − *α* = 99%, assumption that #FP is Poisson-distributed).

Consider the comparison of the red and the brown points labeled *k* = 15 in Figure [3]. Here, M5 (brown) received the seeds output from M3 (red) as input and provided an additional filter. With it, geometric hashing reduced the number of false positives produced by a set of four spaced seed patterns (M3) of weight *k* = 15 from 11.2 false seeds per genome position (#FP = 8,563,355) to arguably less than 6.2 · 10^−6^ false seeds per genome position (#FP = 0), i.e. by a factor of about 2 · 10^6^. At the same time, the sensitivity only decreased slightly from 0.974 to 0.972.

When using a smaller *k*, seed finding with geometric hashing can simultaneously be more sensitive and much more specific than sets of spaced seeds. To see this, compare geometric hashing (M5) for *k* = 14 with M3 (four patterns) for *k* = 15. With this configuration, M5 has 2.1% false negatives and 279 FP while M3 has 2.6% false negatives and 88,563,355 FP (see Figure [3] and Supplementary Material). Stated differently, M5 reduces the false negatives by 0.5%*/*2.6% ≈ 19% and the false positives by a factor ∼ 3 · 10^5^.

M4 (neighbouring matches) is able to achieve zero false positives as well, however at lower sensitivity. Among the baseline methods M1-M3, contiguous seeds have the worst trade-off between sensitivity and 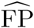. Using one spaced seed patterns greatly improves this trade-off, which is even better when using two or four spaced seed patterns. M4 and M5 were run with four spaced seed patterns and are thus directly comparable to the performance of M3 with four patterns. We compared the runtime of M3, M4 and M5 when all were run with the same spaced seed patterns in Table [1]. M4 and M5 both need only marginally more time to improve the specificity. However, M4 reduces the sensitivity stronger than M5 compared to M3.

**Table 1:** Runtime and memory requirements of comparable runs of M3 (multiple spaced seed patterns), M4 (neighbouring matches) and M5 (geometric hashing).

| Method | weight | patterns | sensitivity | $\widehat{\text{FP}}$ | additional runtime [s] |
| --- | --- | --- | --- | --- | --- |
| M3 | 15 | 4 | 0.974 | 11.2 | – |
| M4 | 15 | 4 | 0.887 | $39 \cdot 10^{-6}$ | 41 |
| M5 | 15 | 4 | 0.972 | 0 | 107 |

### 2.1 Metaparameter Optimization

The key parameter for seed finding is the weight *k* of the seeds. We determined optimal weights minimizing 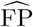 while requiring that the sensitivity is *at least* 0.9. Figure [4] shows the FP count for each method at the optimal weight (numbers over the bars). In Table [2] we compare the performance of the respective runs. Using multiple spaced seeds (M3) alone heavily increases runtime and memory requirement, which can be reduced with smaller weights. Even though geometric hashing is neither the fastest nor requires the least memory, it yields the highest sensitivity as it produced no FPs. Note that we also did not focus on optimizing our code to get the best performance possible and that memory requirements for geometric hashing are not directly comparable to the other methods due to implementation details. We performed grid searches on the remaining metaparameters for the respective methods. Neighbouring matches (M4) was run with a neighbour count threshold *τ* = 2 and a search area *D* = 1000. Geometric hashing was run with a tile size *F* = 10, 000, *b* = 500 sub-tiles, a normalization parameter *η* = 300, 000, 000 and a tile score threshold *τ* = 25.

**Table 2:** Runtime and memory requirements of the methods when run with best weight as determined by Figure [4]. The weights were chosen such that each method had the lowest possible 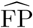 but a sensitivity of at least 0.9. Note that the memory requirement of M5 (geometric hashing) cannot be directly compared to the other methods as of implementation differences.

| Method | weight | patterns | sensitivity | $\widehat{\text{FP}}$ | runtime [min] | memory [GB] |
| --- | --- | --- | --- | --- | --- | --- |
| M1 | 14 | 1 | 0.911 | 11.8 | 8 | 13.3 |
| M2 | 17 | 1 | 0.900 | 0.166 | 6 | 23.0 |
| M3 | 19 | 2 | 0.903 | 0.0155 | 12 | 51.7 |
| M3 | 20 | 4 | 0.914 | 0.0115 | 26 | 122.4 |
| M4 | 14 | 4 | 0.919 | 0.000497 | 26 | 69.5 |
| M5 | 15 | 4 | 0.972 | 0 | 29 | 129.3 |

**Figure 4:**
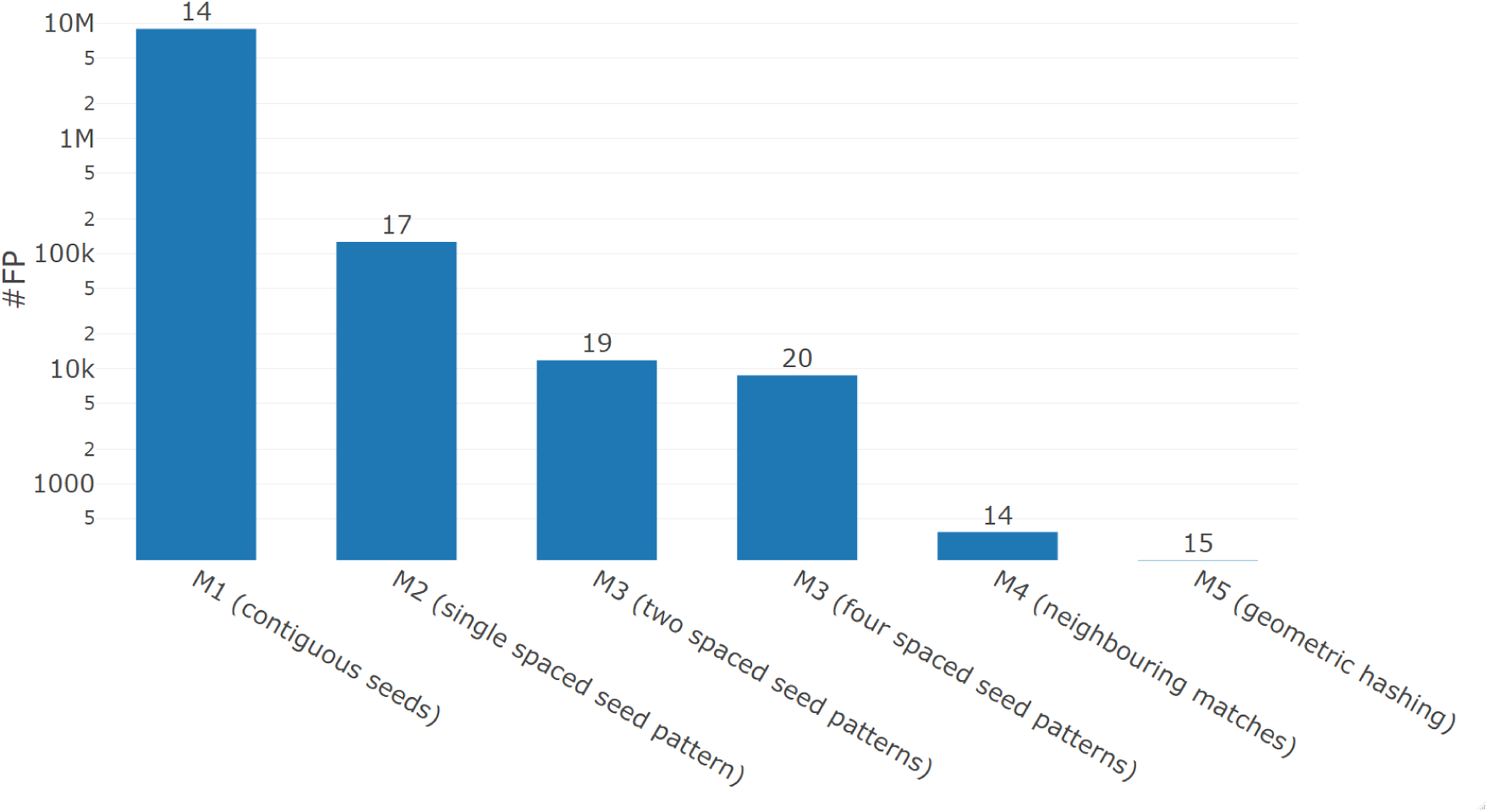
Comparison of minimal achievable FP count when sensitivity is required to be at least 0.9. Geometric hashing is the only method that does not report any false positives in our dataset at this sensitivity threshold. The numbers on top of the bars denote the weight of the seed or spaced seed pattern(s). On the *y*-axis k and M abbreviate 10^3^ and 10^6^, respectively.

## 3 Discussion

Our experiments show that geometric hashing is the most accurate method on our test data. Geometric hashing can reduce the number of false positives by about 6 orders of magnitude over the previous state of the art in seed finding – sets of (four) spaced seed patterns. The later method itself constitutes an improvement of 3-4 orders of magnitude over the naive method of using contiguous *k*-mer matches as alignment seeds.

Geometric hashing can be used as a filter on any other seed finding method and may potentially be ‘inserted’ into the internal pipeline of existing alignment programs between seed finding and seed extension. A large speed-up may be possible in such aligners if significant parts of the runtime are spent downstream of seed finding. Note that seed extension, followed by scoring and filtering, is also an algoritmic step to achieve a higher specificity. However, it requires running an algorithm for local alignment scoring over a variable-length region. In constrast, geometric hashing only requires a constant number of simple integer arithmetic functions (in fact two: one for mapping, one for scoring) and does not require any sequence comparisons. A seed extension phase would still follow but needs to be executed on a much smaller set of seeds.

Moreover, a higher sensitivity can be achieved by lowering *k*. Due to the effective filtering of geometric hashing, this need not come at the computational cost of a large number of false positives. The seed finding with geometric hashing can then simultaneously be somewhat more sensitive and much more specific than sets of spaced seeds. Even small improvements in sensitivity may result in finding orthologies with lower sequence similarity [34].

With our results we confirmed the well established advantage of spaced seeds (M2, M3) over contiguous seeds (M1) and that additional filtering steps (M4, M5) can improve seed performance even further. The focus of this work was to show that considering even distant exons in filtering, as done in geometric hashing, works and is superior to only considering local neighbourhoods of seeds candidates as done in M4. Due to its ability to filter out FP efficiently, we expect a large runtime improvement from geometric hashing when applied in pairwise whole-genome alingments, as much fewer wrong seeds have to be considered in the more time consuming seed extension phase.

We think that the geometric hashing idea is not limited to genome alignments. It could be adapted for protein sequences as well and with some adjustments also for protein to mRNA alignments, or it could be used to speed up tBLASTX [3], which finds region pairs in genomes that are similar peptide sequences when translated. As mentioned earlier, the geometric hashing idea also generalizes well for higher dimensional seeds that occur when multiple genomes are compared simultaneously. Future work could focus on the necessary adaptations for this to work well.

The methods for seed filtering can be seen as proofs of concept with much potential for improvements as we did not aim to write a fully optimized software. For example, it has been reported that sorting matches by keys rather than hashing can improve runtime due to better data locality [22]. Such an implementation is compatible with our geometric hashing approach, both for the primary hashing of *k*-mers and for the seconary geometric hashing of tiles. Further, the overall memory footprint can be decreased when the genomes are processed in chunks rather than all at once.

## 4 Conclusion

We presented a novel seed filtering approach, *geometric hashing*, that uses non-local neighbourhood information to find orthologous genes with high precision. It outperforms local neighbourhood filtering while only slightly affecting the sensitivity of unfiltered seeds. Geometric hashing is a simple yet powerful idea and generalizes well to other alignment problems and higher dimensional data and could be a strong tool in future (multiple) genome alignment approaches.

## Supporting information

Supplementary Material - Raw Accuracy

## Declarations

### Ethics approval and consent to participate

Not applicable

### Consent for publication

Not applicable

### Availability of data and materials

The code and test data is available at our Github repository [41].

## Acknowledgements

We thank Andre Kahles, Gunnar Rätsch, Amir Joudaki, Mikhail Karasikov and Harun Mustafa from the Biomedical Informatics Group at ETH Zürich for regular feedback and discussions.

## Author’s contributions

ME implemented all code, ran all experiments, visualized and interpreted the results and wrote the manuscript. GM prepared the test data and wrote the manuscript. MS conceived the geometric hashing idea, provided academic guidance throughout the project and wrote the manuscript. All authors read and approved the final manuscript.

## References

[1] Armstrong, J., Fiddes, I.T., Diekhans, M., Paten, B.: Whole-genome alignment and comparative annotation. Annual Review of Animal Biosciences 7(1), 41–64 (2019). doi: 10.1146/annurev-animal-020518-115005. PMID: 30379572. https://doi.org/10.1146/annurev-animal-020518-115005

[2] Vertebrate Genomes Project. https://vertebrategenomesproject.org/ [Accessed: 2020-2-21] (2020)

[3] Altschul, S.F., Gish, W., Miller, W., Myers, E.W., Lipman, D.J.: A basic local alignment search tool. J Mol Biol 215, 403–410 (1990)

[4] Ma, B., Tromp, J., Li, M.: PatternHunter: faster and more sensitive homology search. Bioinformatics 18(3), 440–445 (2002)

[5] Burkhardt, S., Kärkkäinen, J.: Better filtering with gapped q-grams. Fundamenta informaticae 56(1-2), 51–70 (2003)

[6] Li, M., Ma, B., Kisman, D., Tromp, J.: Patternhunter ii: Highly sensitive and fast homology search. Journal of bioinformatics and computational biology 2(03), 417–439 (2004)

[7] Keich, U., Li, M., Ma, B., Tromp, J.: On spaced seeds for similarity search. Discrete Applied Mathematics 138(3), 253–263 (2004). doi: 10.1016/S0166-218X(03)00382-2

[8] Choi, K.P., Zeng, F., Zhang, L.: Good spaced seeds for homology search. Bioinformatics 20(7), 1053–1059 (2004). doi: 10.1093/bioinformatics/bth037. https://academic.oup.com/bioinformatics/article-pdf/20/7/1053/678172/bth037.pdf

[9] Choi, K.P., Zhang, L.: Sensitivity analysis and efficient method for identifying optimal spaced seeds. Journal of Computer and System Sciences 68(1), 22–40 (2004). doi: 10.1016/j.jcss.2003.04.002

[10] Brejová, B., Brown, D.G., Vinař, T.: Optimal spaced seeds for homologous coding regions. Journal of Bioinformatics and Computational Biology 01(04), 595–610 (2004). doi: 10.1142/S0219720004000326. https://doi.org/10.1142/S0219720004000326

[11] Buhler, J., Keich, U., Sun, Y.: Designing seeds for similarity search in genomic DNA. Journal of Computer and System Sciences 70(3), 342–363 (2005). doi: 10.1016/j.jcss.2004.12.003. Special Issue on Bioinformatics II

[12] Sun, Y., Buhler, J.: Designing multiple simultaneous seeds for DNA similarity search. Journal of computational biology : a journal of computational molecular cell biology 12(6), 847–861 (2005). doi: 10.1089/cmb.2005.12.847

[13] Li, M., Ma, B., Zhang, L.: Superiority and complexity of the spaced seeds. In: Proceedings of the Seventeenth Annual ACM-SIAM Symposium on Discrete Algorithm. SODA’06, pp. 444–453. Society for Industrial and Applied Mathematics, USA (2006)

[14] Kucherov, G., Noe, L., Ponty, Y.: Estimating seed sensitivity on homogeneous alignments. In: Proceedings. Fourth IEEE Symposium on Bioinformatics and Bioengineering, pp. 387–394 (2004). doi: 10.1109/BIBE.2004.1317369

[15] Ma, B., Li, M.: On the complexity of the spaced seeds. Journal of Computer and System Sciences 73(7), 1024–1034 (2007). doi: 10.1016/j.jcss.2007.03.008. Bioinformatics III

[16] Nicolas, F., Rivals, E.: Hardness of optimal spaced seed design. Journal of Computer and System Sciences 74(5), 831–849 (2008). doi: 10.1016/j.jcss.2007.10.001

[17] Ilie, L., Ilie, S.: Multiple spaced seeds for homology search. Bioinformatics 23(22), 2969–2977 (2007). doi: 10.1093/bioinformatics/btm422. http://oup.prod.sis.lan/bioinformatics/article-pdf/23/22/2969/543804/btm422.pdf

[18] Farach-Colton, M., Landau, G.M., Sahinalp, S.C., Tsur, D.: Optimal spaced seeds for faster approximate string matching. Journal of Computer and System Sciences 73(7), 1035–1044 (2007). doi: 10.1016/j.jcss.2007.03.007. Bioinformatics III

[19] Mak, D.Y.F., Benson, G.: All hits all the time: parameter-free calculation of spaced seed sensitivity. Bioinformatics 25(3), 302–308 (2008). doi: 10.1093/bioinformatics/btn643. https://academic.oup.com/bioinformatics/article-pdf/25/3/302/675637/btn643.pdf

[20] Chung, W.-H., Park, S.-B.: Hit integration for identifying optimal spaced seeds. BMC Bioinformatics 11(1), 37 (2010). doi: 10.1186/1471-2105-11-S1-S37

[21] Brown, D.G.: A survey of seeding for sequence alignment. Bioinformatics algorithms: techniques and applications, 117–142 (2008)

[22] Buchfink, B., Xie, C., Huson, D.H.: Fast and sensitive protein alignment using diamond. Nature Methods 12(1), 59–60 (2015). doi: 10.1038/nmeth.3176

[23] Harris, R.S.: Improved pairwise alignment of genomic DNA. PhD thesis, Pennsylvania State University (2007)

[24] Noé, L., Kucherov, G.: Yass: enhancing the sensitivity of DNA similarity search. Nucleic Acids Research 33(suppl 2), 540–543 (2005). doi: 10.1093/nar/gki478. https://academic.oup.com/nar/article-pdf/33/suppl2/W540/7623590/gki478.pdf

[25] NCBI: BLAST topics. https://blast.ncbi.nlm.nih.gov/Blast.cgi?CMD=Web&PAGE_TYPE=BlastDocs&DOC_TYPE=BlastHelp#discMegaBlast [Accessed: 2020-02-12] (2020)

[26] Morgulis, A., Coulouris, G., Raytselis, Y., Madden, T.L., Agarwala, R., Schäffer, A.A.: Database indexing for production MegaBLAST searches. Bioinformatics (Oxford, England) 24(16), 1757–1764 (2008). doi: 10.1093/bioinformatics/btn322. 18567917[pmid]

[27] Noé, L., Kucherov, G.: Improved hit criteria for DNA local alignment. BMC Bioinformatics 5(1), 149 (2004). doi: 10.1186/1471-2105-5-149

[28] Mak, D., Gelfand, Y., Benson, G.: Indel seeds for homology search. Bioinformatics 22(14), 341–349 (2006). doi: 10.1093/bioinformatics/btl263. https://academic.oup.com/bioinformatics/article-pdf/22/14/e341/617355/btl263.pdf

[29] Leimeister, C.-A., Dencker, T., Morgenstern, B.: Accurate multiple alignment of distantly related genome sequences using filtered spaced word matches as anchor points. Bioinformatics 35(2), 211–218 (2018)

[30] GRCh38.p13 [Internet]. Bethesda (MD): National Library of Medicine (US), National Center for Biotechnology Information. [cited 2020 02 20]. Available from: https://www.ncbi.nlm.nih.gov/assembly/ (2012)

[31] GRCm38.p6 [Internet]. Bethesda (MD): National Library of Medicine (US), National Center for Biotechnology Information. [cited 2020 02 20]. Available from: https://www.ncbi.nlm.nih.gov/assembly/ (2012)

[32] Kinsella, R.J., Kähäri, A., Haider, S., Zamora, J., Proctor, G., Spudich, G., Almeida-King, J., Staines, D., Derwent, P., Kerhornou, A., Kersey, P., Flicek, P.: Ensembl BioMarts: a hub for data retrieval across taxonomic space. Database 2011 (2011). doi: 10.1093/database/bar030. bar030. https://academic.oup.com/database/article-pdf/doi/10.1093/database/bar030/1262975/bar030.pdf

[33] Cunningham, F., Achuthan, P., Akanni, W., Allen, J., Amode, M., Armean, I.M., Bennett, R., Bhai, J., Billis, K., Boddu, S., Cummins, C., Davidson, C., Dodiya, K.J., Gall, A., Girón, C.G., Gil, L., Grego, T., Haggerty, L., Haskell, E., Hourlier, T., Izuogu, O.G., Janacek, S.H., Juettemann, T., Kay, M., Laird, M.R., Lavidas, I., Liu, Z., Loveland, J., Marugán, J.C., Maurel, T., McMahon, A.C., Moore, B., Morales, J., Mudge, J.M., Nuhn, M., Ogeh, D., Parker, A., Parton, A., Patricio, M., Abdul Salam, A.I., Schmitt, B.M., Schuilenburg, H., Sheppard, D., Sparrow, H., Stapleton, E., Szuba, M., Taylor, K., Threadgold, G., Thormann, A., Vullo, A., Walts, B., Winterbottom, A., Zadissa, A., Chakiachvili, M., Frankish, A., Hunt, S.E., Kostadima, M., Langridge, N., Martin, F.J., Muffato, M., Perry, E., Ruffier, M., Staines, D.M., Trevanion, S.J., Aken, B.L., Yates, A.D., Zerbino, D.R., Flicek, P.: Ensembl 2019. Nucleic Acids Research 47(D1), 745–751 (2018). doi: 10.1093/nar/gky1113. https://academic.oup.com/nar/article-pdf/47/D1/D745/27437646/gky1113.pdf

[34] Hahn, L., Leimeister, C.-A., Ounit, R., Lonardi, S., Morgenstern, B.: Rasbhari: optimizing spaced seeds for database searching, read mapping and alignment-free sequence comparison. PLoS Computational Biology 12(10), 1005107 (2016)

[35] Ilie, L., Ilie, S., Mansouri Bigvand, A.: SpEED: fast computation of sensitive spaced seeds. Bioinformatics 27(17), 2433–2434 (2011). doi: 10.1093/bioinformatics/btr368. http://oup.prod.sis.lan/bioinformatics/article-pdf/27/17/2433/16898964/btr368.pdf

[36] Ilie, L., Ilie, S.: Long spaced seeds for finding similarities between biological sequences. BIOCOMP 7, 25–28 (2007)

[37] Makalowski, W., Zhang, J., Boguski, M.S.: Comparative analysis of 1196 orthologous mouse and human full-length mRNA and protein sequences. Genome Research 6(9), 846–857 (1996)

[38] Wolfson, H.J., Rigoutsos, I.: Geometric hashing: an overview. IEEE Computational Science and Engineering 4(4), 10–21 (1997). doi: 10.1109/99.641604

[39] Kent, W.J., Sugnet, C.W., Furey, T.S., Roskin, K.M., Pringle, T.H., Zahler, A.M., Haussler, D.: The Human Genome Browser at UCSC. Genome Research 12(6), 996–1006 (2002). doi: 10.1101/gr.229102. Article published online before print in May 2002. http://www.genome.org/cgi/reprint/12/6/996.pdf

[40] UCSC Genome Browser. http://genome.ucsc.edu/ [Accessed: 2020-2-20] (2020)

[41] Geometric Hashing. Github. https://github.com/Gaius-Augustus/GeometricHashing [cited 2020-02-21] (2020)

