## Supplementary Material - Raw Accuracy for "Global, Highly Specific and Fast Filtering of Alignment Seeds"

May 1, 2020

### 1 M4: Neighbouring Matches Algorithm

For this method, the seed candidates $(S_1, i, S_2, j)$ are sorted. The primary sort criterion is the sequence pair $(S_1, S_2)$, the secondary sort criterion is the diagonal $i - j$, the tertiary sort criterion is the position $i$. This sorting is achieved in $O(n \log(n))$ time where $n$ is the number of seed candidates. In line 3 of below algorithm, GetNextDiagonal() returns a list of seeds that all lie on the same diagonal $i - j$, one diagonal at a time. This is linear in the length $d$ of the respective diagonal. The body of the loop in line 4 consumes each seed exactly once and is in total executed $n$ times. The check in line 5 can be implemented $O(1)$ amortized time.

Neighbouring matches filter algorithm

```
1: Sort(S) // sort set S of seed candidates
2: repeat
3:     S′ = GetNextDiagonal(S)
4:     for all s ∈ S′ do
5:         if numNeighbors(s, S′) ≥ τ then
6:             report s
7: until all diagonals processed
```

### 2 Raw Accuracy Data

Tables [1], [2], [3], [4] and [5] list the values plottet for each method in Figure 3 of the main text.

**Supplementary Table 1:** Details for M1 (contiguous seeds) for different weights

| weight | patterns | sensitivity | $\widehat{\mathrm{FP}}$ | #FP |
|---|---|---|---|---|
| 13 | 1 | 0.928 | 44.7 | 34165813 |
| 14 | 1 | 0.911 | 11.8 | 8984036 |
| 15 | 1 | 0.853 | 2.91 | 2228492 |
| 16 | 1 | 0.825 | 0.704 | 538566 |
| 17 | 1 | 0.798 | 0.169 | 128859 |
| 18 | 1 | 0.729 | 0.0368 | 28116 |
| 19 | 1 | 0.698 | 0.00770 | 5891 |
| 20 | 1 | 0.670 | 0.00203 | 1552 |
| 21 | 1 | 0.593 | 0.000637 | 487 |
| 22 | 1 | 0.565 | 0.000158 | 121 |
| 23 | 1 | 0.536 | 0.0000131 | 10 |
| 24 | 1 | 0.468 | 0.00000131 | 1 |

**Supplementary Table 2:** Details for M2 (single spaced seed pattern) for different weights

| weight | patterns | sensitivity | $\widehat{\mathrm{FP}}$ | #FP |
|---|---|---|---|---|
| 13 | 1 | 0.967 | 45.2 | 34587566 |
| 14 | 1 | 0.957 | 11.3 | 8639856 |
| 15 | 1 | 0.936 | 2.76 | 2112892 |
| 16 | 1 | 0.920 | 0.712 | 544387 |
| 17 | 1 | 0.900 | 0.166 | 126779 |
| 18 | 1 | 0.868 | 0.0350 | 26731 |
| 19 | 1 | 0.849 | 0.0156 | 11949 |
| 20 | 1 | 0.828 | 0.00182 | 1394 |
| 21 | 1 | 0.805 | 0.000649 | 496 |
| 22 | 1 | 0.757 | 0.0000379 | 29 |
| 23 | 1 | 0.745 | 0.0000144 | 11 |
| 24 | 1 | 0.714 | 0.00000131 | 1 |

**Supplementary Table 3:** Details for M3 (multiple spaced seed patterns) for different weights

| weight | patterns | sensitivity | $\widehat{\text{FP}}$ | #FP |
|---|---|---|---|---|
| 13 | 2 | 0.981 | 89.5 | 68412504 |
| 14 | 2 | 0.974 | 23.6 | 18017544 |
| 15 | 2 | 0.965 | 5.93 | 4530575 |
| 16 | 2 | 0.951 | 1.48 | 1131934 |
| 17 | 2 | 0.936 | 0.351 | 268548 |
| 18 | 2 | 0.915 | 0.0803 | 61395 |
| 19 | 2 | 0.903 | 0.0155 | 11828 |
| 20 | 2 | 0.887 | 0.00690 | 5276 |
| 21 | 2 | 0.872 | 0.000815 | 623 |
| 22 | 2 | 0.838 | 0.0000562 | 43 |
| 23 | 2 | 0.792 | 0.0000314 | 24 |
| 24 | 2 | 0.771 | 0.00000916 | 7 |
| 13 | 4 | 0.987 | 179 | 136648882 |
| 14 | 4 | 0.982 | 45.9 | 35090192 |
| 15 | 4 | 0.974 | 11.2 | 8563355 |
| 16 | 4 | 0.965 | 2.84 | 2167649 |
| 17 | 4 | 0.956 | 0.678 | 518284 |
| 18 | 4 | 0.945 | 0.156 | 119128 |
| 19 | 4 | 0.929 | 0.0446 | 34091 |
| 20 | 4 | 0.914 | 0.0115 | 8766 |
| 21 | 4 | 0.900 | 0.00229 | 1754 |
| 22 | 4 | 0.879 | 0.000511 | 391 |
| 23 | 4 | 0.874 | 0.000233 | 178 |
| 24 | 4 | 0.826 | 0.0000419 | 32 |

**Supplementary Table 4:** Details for M4 (neighbouring matches) for different weights

| weight | patterns | sensitivity | $\widehat{\text{FP}}$ | #FP |
|---|---|---|---|---|
| 13 | 4 | 0.932 | 0.00974 | 7444 |
| 14 | 4 | 0.919 | 0.000497 | 380 |
| 15 | 4 | 0.887 | 0.0000392 | 30 |
| 16 | 4 | 0.857 | 0.00000262 | 2 |
| 17 | 4 | 0.819 | 0 | 0 |
| 18 | 4 | 0.771 | 0 | 0 |
| 19 | 4 | 0.732 | 0 | 0 |
| 20 | 4 | 0.682 | 0 | 0 |
| 21 | 4 | 0.645 | 0 | 0 |
| 22 | 4 | 0.606 | 0 | 0 |
| 23 | 4 | 0.602 | 0 | 0 |
| 24 | 4 | 0.512 | 0 | 0 |

**Supplementary Table 5:** Details for M5 (geometric hashing) for different weights

| weight | patterns | sensitivity | $\widehat{\text{FP}}$ | #FP |
|---|---|---|---|---|
| 13 | 4 | 0.984 | 14.9 | 11359790 |
| 14 | 4 | 0.979 | 0.000365 | 279 |
| 15 | 4 | 0.972 | 0 | 0 |
| 16 | 4 | 0.963 | 0 | 0 |
| 17 | 4 | 0.952 | 0 | 0 |
| 18 | 4 | 0.939 | 0 | 0 |
| 19 | 4 | 0.922 | 0 | 0 |
| 20 | 4 | 0.905 | 0 | 0 |
| 21 | 4 | 0.887 | 0 | 0 |
| 22 | 4 | 0.864 | 0 | 0 |
| 23 | 4 | 0.858 | 0 | 0 |
| 24 | 4 | 0.803 | 0 | 0 |
